# Magainin 2 and PGLa in Bacterial Membrane Mimics II: Membrane Fusion and Sponge Phase Formation

**DOI:** 10.1101/763383

**Authors:** Ivo Kabelka, Michael Pachler, Sylvain Prévost, Ilse Letofsky-Papst, Karl Lohner, Georg Pabst, Robert Vácha

## Abstract

We studied the synergistic mechanism of equimolar mixtures of magainin 2 (MG2a) and PGLa in phosphatidylethanolamine/phosphatidylglycerol mimics of Gram-negative cytoplasmic membranes. In a preceding paper [Pachler et al., Biophys. J. 2019 xxx], we reported on the early onset of parallel heterodimer formation of the two antimicrobial peptides already at low concentrations and the resulting defect formation in membranes. Here, we focus on the structures of the peptide/lipid aggregates occurring in the synergistic regime at elevated peptide concentrations. Using a combination of calorimetric, scattering, electron microscopic and *in silico* techniques, we demonstrate that the two peptides, even if applied individually, transform originally large unilamellar vesicles into multilamellar vesicles, with a collapsed interbilayer spacing resulting from peptide induced adhesion. Interestingly, the adhesion does not lead to a peptide induced lipid separation of charged and charge neutral species. In addition to this behavior, equimolar mixtures of MG2a and PGLa formed surface-aligned fibril-like structures, which induced adhesion zones between the membranes and the formation of transient fusion stalks in molecular dynamics simulations and a coexisting sponge phase observed by small-angle X-ray scattering. The previously reported increased leakage of lipid vesicles of identical composition in the presence of MG2a/PGLa mixtures is therefore related to a peptide-induced cross-linking of bilayers.

**STATEMENT OF SIGNIFICANCE:** We demonstrate that the synergistic activity of the antimicrobial peptides MG2a and PGLa correlates to the formation of surface-aligned fibril-like peptide aggregates, which cause membrane adhesion, fusion and finally the formation of a sponge phase.

## INTRODUCTION

PGLa and Magainin 2 (MG2a) are two antimicrocial peptides (AMPs) derived from the African clawed frog, which are well known for their synergistic killing of bacteria (1). So far, no consensus has been reached regarding the biophysical mechanism though. While some groups proposed the formation of distinct transmembrane pores formed by PGLa/MG2a dimers (1) or complex peptide aggregates (2), others suggested the formation of mesophase-like structures by surface-aligned peptides (3). Recently, we reported that the activity of these peptides in *Escherichia coli* can be reasonably mimicked by mixtures of palmitoyl oleoyl phosphatidylethanolamine (POPE) and palmitoyl oleoyl phosphatidylglycerol (POPG) (molar ratio 3:1), while other lipid mixtures fail to capture synergistic effects (4). Intriguingly, both peptides, in particular when mixed at an equimolar ratio, were shown to adopt a surface-aligned topology in POPE-enriched bilayers (5, 6), thus questioning the formation of transmembrane pores. However, even if the peptides from mesophase-like structures at synergistic concentrations in POPE-enriched bilayers (3), it is far from being clear how that should cause enhanced leakage of dyes from large unilamellar vesicles (LUVs) (4). In order to address the synergistic mechanism, we applied several experimental and computational tools that allowed us to study the peptide-induced effects occurring on the nanoscopic to macroscopic length scales. In the preceding paper of this series, we combined small-angle X-ray and neutron scattering (SAXS/SANS) experiments with all-atom and coarse-grained molecular dynamics simulations to show that equimolar mixtures of L18W-PGLa and MG2a are prone to form surface-aligned parallel heterodimers already at low peptide concentrations and the dimerisation leads to an increase of the defects within the membrane interior (7).

The present paper focuses on the synergistic regime of the peptides at high concentrations (i.e. peptide to lipid molar ratio P/L ≤ 1/ 50) (4). For direct comparison to our previous results including leakage assays, we used L18W-PGLa instead of native PGLa (4). Nevertheless, the mutant behavior was reported to be analogous to native PGLa (1). In addition to the experimental techniques applied in our first paper (7), we used differential scanning calorimetry (DSC) to reveal effects of the peptides on the cooperative melting of POPE/POPG bilayers, as well as cryogenic transmission electron microscopy (cryo-TEM) to capture the macroscopic equilibrium structure after incubating LUVs with the two peptides. Our results demonstrate that the peptides transform the LUVs at sufficiently high concentration into condensed multilamellar vesicles (MLVs) with the peptides sandwiched between the individual bilayers. Additionally, equimolar peptide mixtures formed surface-aligned fibril-like structures in the interstitial bilayer space, which finally lead to fusion events and to the formation of a leaky sponge phase.

## MATERIALS AND METHODS

### Experiments

#### Lipids, Peptids and Chemicals

POPE, POPG, palmitoyl-d31-oleoyl-phosphoethanolamine (POPE-d31) and palmitoyl-d31-oleoyl-phosphatidylglycerol (POPG-d31) were purchased from Avanti Polar Lipids (Alabaster, AL, USA, purity > 99%) as powder. L18W-PGLa, MG2a and the chemically linked heterodimer L18W-PGLa–MG2a, denoted in the following as hybrid peptide (Fig. S1), were obtained in lyophilized form (purity >95%) from PolyPeptide Laboratories (San Diego, CA, USA). Deuterium oxide (D_2_O, purity 99.8 atom%) and HEPES (purity >99.5) were purchased from Carl Roth (Karl-sruhe, Baden-Wüttenberg, Germany). All other chemicals were obtained from Sigma-Aldrich (Vienna, Austria) in *pro analysis* quality. Lipid stock solutions for sample preparation were prepared in organic solvent chloroform/methanol (9:1; v/v) and phosphate assayed for quantification of lipid content (8). Peptide stock solutions were prepared in 10 mM HEPES 140 mM NaCl buffer solution (pH 7.4).

#### Sample preparation

Samples were prepared, assayed and controlled as described in (7). Samples for zero-contrast neutron experiments contained mixtures of protiated and deuterated lipids as detailed below and were hydrated in 10 mM HEPES (140 mM NaCl, pH 7.4) prepared in D_2_O/H_2_O (43.7 vol%). We emphasize that all lipid samples were extruded to ∼100 nm-sized large unilamellar vesicles (LUVs) prior to the addition of the peptides. Further, all presented data have been recorded at least three days after peptide addition. We therefore refer to these data as end states.

### Small-Angle X-ray Scattering (SAXS)

SAXS data were collected at the SWING beamline (Soleil, Saint-Aubin, France) as described in (7). Peptide induced aggregation of vesicles led to the formation of Bragg peaks, which impeded the data analysis developed in (7). Instead, data were analyzed based on Bragg peak positions and intensities only, see e.g. (9). In brief, the electron density for a centrosymetric lamellar structure is given by

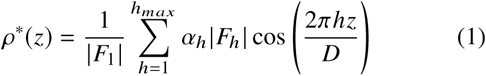

where *α*_*h*_ = ± 1 is the phase, *F*_*h*_ is the form factor of the h’ reflection, and *D* = 2*πh* / *q*_*h*_ is lamellar repeat distance. From the derived electron density profiles we calculated the head to headgroup distance *D*_HH_ from the distance between the maxima across the lipid bilayer. The steric bilayer thickness *D ′*_*B*_ = *D*_HH_ + 2*D*_PO_ can be estimated by using *D*_PO_ = 2.92 Å, which corresponds to a distance between the PO_4_ group and the furthest outward lipid group (here the glycerol group) as determined from a joint SAXS/SANS data analysis (7).

Bragg peak positions of cubic phases index as

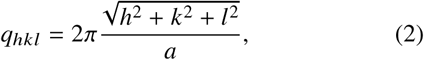

where *h, k, l* are the the Miller indices and *a* is the lattice parameter.

### Small-Angle Neutron Scattering (SANS)

Zero-contrast SANS data were acquired on D11 at the Institut Laue-Langevin (ILL), Grenoble, France, with a multiwire ^3^He detector of 256 × 256 pixels (3.75 × 3.75 mm^2^), at a wave length *λ* = 0.53 nm (Δ *λ* / *λ* = 9 %), and three sample-to-detector distances: 1.4, 8 and 39 m (with corresponding collimations of 8 m, 8 m and 40.5 m, respectively, covering a *q*-range of 0.015 5 - nm^-1^. Samples were poured into quartz cuvettes of either 2 mm or 1 mm pathway and mounted in a thermostated rotating holder to minimize sample sedimentation. Data were reduced with BerSANS, accounting for flat field, solid angle, dead time, transmission, and background subtraction. Scattered intensities were placed on an absolute scale using the scattering by 1 mm H_2_O as a secondary standard.

In general, the scattered intensity is proportional to the scattering contrast Δ *ρ*

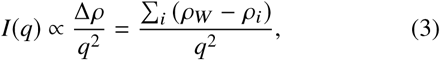

where *ρ*_*W*_ is the neutron scattering length density (NSLD) of the buffer, and *ρ*_*i*_ is the NSLD of a given slab of the lipid bilayer. For designing zero contrast conditions, we considered two slabs describing the lipid headgroup and hydrocarbon core, respectively. In this case it can be estimated that homogeneously distributed lipids match the 43.7 vol% deuterated buffer at a molar ratio of 78/22 deuterated/protiated hydrocarbons. That is, under these experimental conditions, the lipid headgroup and the hydrocarbon core are both fully contrast-matched and invisible to neutrons.

We used two lipid mixtures, both having a total deuterated/protiated hydrocarbon ratio of 78/22. System A with POPE-d31/ (POPG)/(POPG-d31) at (75)/(22 /3) mol/mol/mol and system B with (POPE/POPE-d31)/ (POPG/POPG-d31) at (16.5/58.5)/(5.5/19.5) mol/mol/mol/mol. The compositions were designed that a peptide-induced lipid separation will give rise lateral contrast and hence lead to detectable coherent scattering signal for the first mixture only. Analogous experiments have been reported for the formation of nanoscopic lipid domains (10). The second mixture was chosen to contrast-match each lipid component individually. Hence, no scattering contrast can be achieved upon lipid demixing. These samples, therefore served as negative controls.

### Differential Scanning Calorimetry (DSC)

DSC experiments were carried out on a MicroCal VP high-sensitivity DSC (Microcal, Northhampton, MA, USA) at a scan rate of 30 °C/h in the temperature regime of 5 to 60°C. Each experiment consisted of three subsequent heating and cooling scans. Baseline subtraction was performed using Origin (OriginLab, Northampton, MA, USA).

### Cryo Transmission Electron Microscopy (TEM)

All cryo-TEM images were recorded with a Gatan system mounted on a Tecnai12 electron microscope (FEI Company, Hillsboro, OR), equipped with a LaB_6_ filament as described previously (11).

### Molecular dynamics simulations

Molecular dynamics (MD) simulations were performed using GROMACS version 2016.2 (12, 13).

#### All-atom simulations

##### Simulation Settings

POPE and POPG lipids were described using the Slipids force field (14, 15), while MG2a (GIGKFLHSAKKFGKAFVGEIMNS-NH_2_) and L18W-PGLa (GMASKAGAIAGKIAKVAWKAL-NH_2_) were simulated with Amberff99SB-ILDN force field (16, 17). The time integration was performed using the leap-frog algorithm with the time step set to 2 fs. A Nosé-Hoover thermostat (18–20) with a coupling constant of 0.5 ps was used to maintain the temperature at 308.15 K. To ensure proper heating of the system, two coupling groups for peptide+lipid and water+ion atoms were used. A Parrinello-Rahman barostat (21, 22) with a semi-isotropic coupling scheme and a coupling constant of 2 ps was employed for keeping the membrane tensionless and the system at a constant pressure of 1 bar. 3D periodic boundary conditions were applied. Long-ranged electrostatic interactions were treated with the particle mesh Ewald method (23) using the real-space cutoff to 1.2 nm. Lennard-Jones interactions were cutoff at 1.2 nm. All bonds were constrained using the LINCS algorithm. Long-range dispersion corrections were applied for both energy and pressure. (24) For equilibration, the protein backbone atoms were held by positional restraints during a 120 ns simulation of the system. For the next 180 ns long equilibration step, backbone dihedral angles were constrained to maintain a helical structure while letting the peptide free to move.

##### Membrane Stack system

The equlibrated bilayer with parallel MG2a+P18W-PGLa heterodimers in each membrane leaflet was duplicated and shifted by 5 nm in the Z-axis (direction of membrane normal). Each bilayer was composed of 192 POPE and 64 POPG lipids equally distributed in both leaflets. The initial box dimensions were 8.9×8.9×8.5 nm. This system with two stacked membranes was solvated and energy minimized using the steepest descent algorithm. Production MD was performed without any restraints for 1 *µ*s.

##### Peptide Fiber system

A smaller membrane patch composed of 96 POPE and 32 POPG lipids (with initial dimensions of 6.9 × 6.9 × 10 nm) was prepared. Two parallel MG2a+L18W-PGLA heterodimers were inserted into the membrane center in parallel orientation with respect to the membrane plane. The dimers were placed in a fibril with the N-terminus of L18W-PGLa peptide (of the first dimer) interacting with C-terminus of MG2a (of the second dimer), see Fig. S8 *A, B*. Thus, the peptides formed an “infinite” fiber over periodic boundary conditions. The production MD was performed without any restraints for 330 ns.

#### Coarse-grained Simulations

##### Simulation Settings

We employed coarse-grained Martini 2.2 (25–27) force field with four-to-one atom mapping in combination with the leap-frog algorithm with an integration time step of 20 fs. A velocity-rescaling thermostat (modified with a stochastic term) (28) with a coupling constant of 1.0 ps maintained the temperature at 310 K. Protein-lipid and solvent beads were coupled to separate baths to ensure correct temperature distribution. The pressure was kept at 1 bar using the Parrinello-Rahman barostat (21, 22) with a semi-isotropic coupling scheme with coupling constant of 12 ps. All non-bonded interactions were cut off at 1.1 nm and the van der Waals potential was shifted to zero. The relative dielectric constant was set to 15.

As a consequence of coarse-graining, the Martini model does not explicitly describe backbone hydrogen bonds. Thus, the secondary structure has to be imposed on the peptides and maintained throughout the simulation. Both MG2a and L18W-PGLa peptides were modeled as *α*-helices. The peptide C-terminal capping was modeled by removing the charge and changing the backbone bead type to neutral.

##### System Preparation

The lipid membranes were assembled in the XY plane using the CHARMM-GUI web server. (29) All Lipid bilayers were formed from POPE and POPG lipids (3:1 mol:mol), equally distributed in both leaflets.

The following systems with two stacked membranes were prepared: (i) a membrane-only system with two 504-lipid bilayers, (ii) two 504-lipid bilayers containing MG2a and L18W-PGLa (1:1 ratio; P/L = 1:42), (iii) same as (ii) but with P/L = 1:21, (iv) two 2016-lipid bilayers containing MG2a and L18W-PGLa (P/L = 1:21), (v) same as (ii) but with but with the peptides placed only in one leaflet of each bilayer. Roughly four solvent beads were added per lipid and NaCl ions were added at 0.13 M concentration with an excess of sodium ions to neutralize the system net charge. In the coarse-grained representation, one water bead corresponds to four water molecule and ion beads are considered to implicitly contain its first hydration shell. The production dynamics simulation was performed for 20 *µ*s with the exception of system (iv) with the simulation length of 100 *µ*s.

## RESULTS

### Experiments

#### Peptides do not induce lipid domain formation, but distribute non-homogeneously

We first investigated the effects of the peptides on the thermotropic behavior of POPE/POPG (3:1 mol/mol) bilayers using DSC. The pure bilayer exhibited a broad, but single main transition at ∼ 23°C, consistent with a previous report (30), and a hysteresis of Δ *T*_*m*_ ∼ - 2°C, upon cooling (Fig. 1).

**Figure 1:**
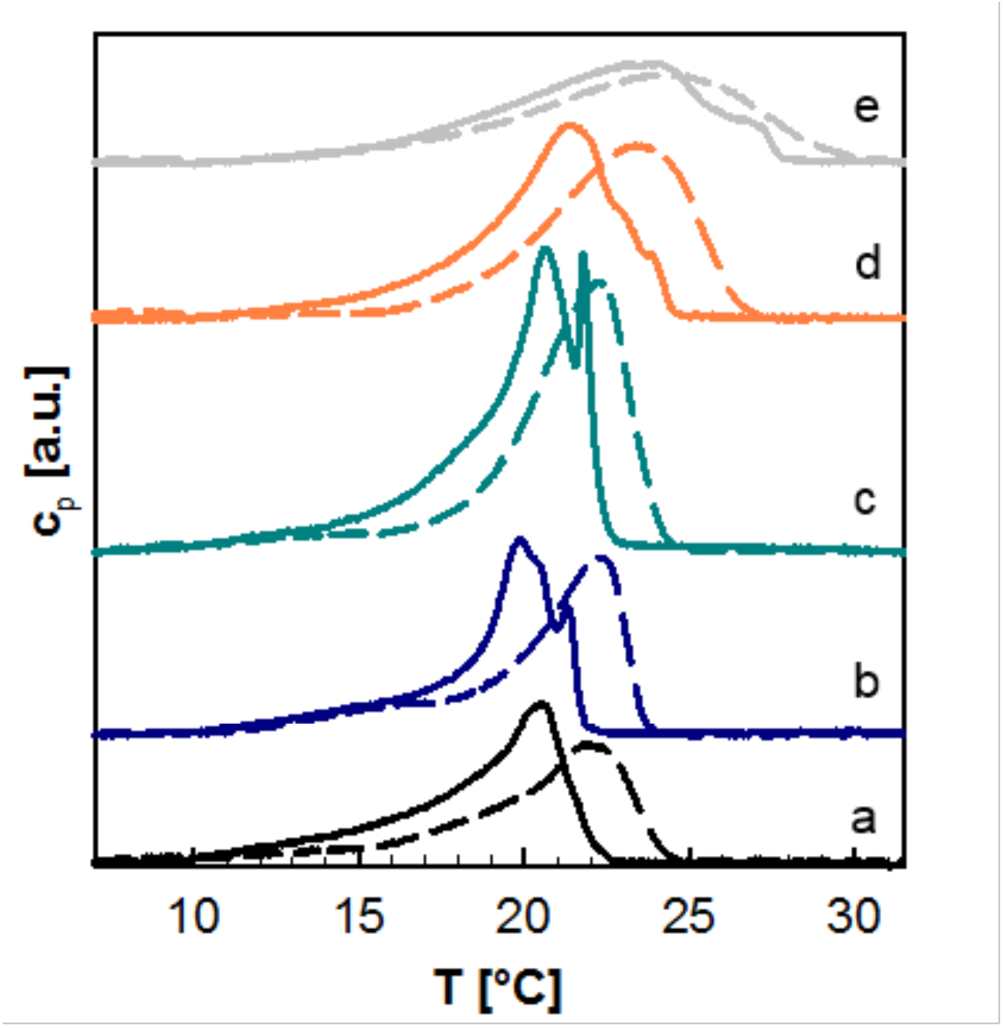
Melting of POPE/POPG (3:1 mol/mol) in the absence and presence of the peptides MG2a (b), L18W-PGLa (c), their equimolar mixture M11 (d) and the hybrid peptide (e) at *P*/ *L* = 1 : 50. Heating scans are shown as dashed lines and cooling scans as solid lines, respectively.

Equilibration of the LUVs in the presence of the peptides (P/L = 1:50) caused a splitting of the melting transition into at least two components, which was mostly pronounced upon cooling. Note that this peptide concentration corresponds to the synergistic regime of both L18W-PGLa/MG2a equimolar mixtures and their chemically-linked hybrid (4). In the case of the equimolar L18W-PGLa/MG2a mixture, denoted in the following as M11, and the hybrid peptide, the splitting of the transition upon cooling occurred over a larger temperature range – and in particular for the hybrid peptide – signifying a loss of cooperativity. Additionally, the maximum of the specific heat capacity shifted slightly to higher temperatures.

In order to test, whether the peptide-induced splitting of the melting transition is due to the formation of lipid domains, we performed zero-contrast SANS experiments. We prepared two lipid mixtures using a combination of perdeuterated and protiated POPE and POPG, which should give rise to a scattering signal in the case of lipid domain formation (system A). System B severed as a negative control and should not show any scattering features even in the case of domain formation (see Material and Methods for details).

Figure 2 shows the corresponding scattering data. Both systems (A and B) exhibited a flat scattering signal for *q* > 0.1 nm^-1^ and increased intensities at low *q*-vectors. The control experiment showed the same trend (Fig. 2 *B*). The increase at low *q*-values can be explained by residual scattering contrast, i.e. slightly imperfect contrast matching, which leads to an increase of scattered intensities due its dependency on *q*^-2^ (see Eq. 3). Additional experiments in the main phase transition regime at P/L = 1:50 yielded analogous results (Figs. S4 - S6). Therefore, we conclude that the addition of peptides does not induce the formation of domains enriched in either lipid species. Consequently, the complex thermotropic behavior revealed by DSC (Fig. 1) needs to be related to a non-homogeneous distribution of peptides.

**Figure 2:**
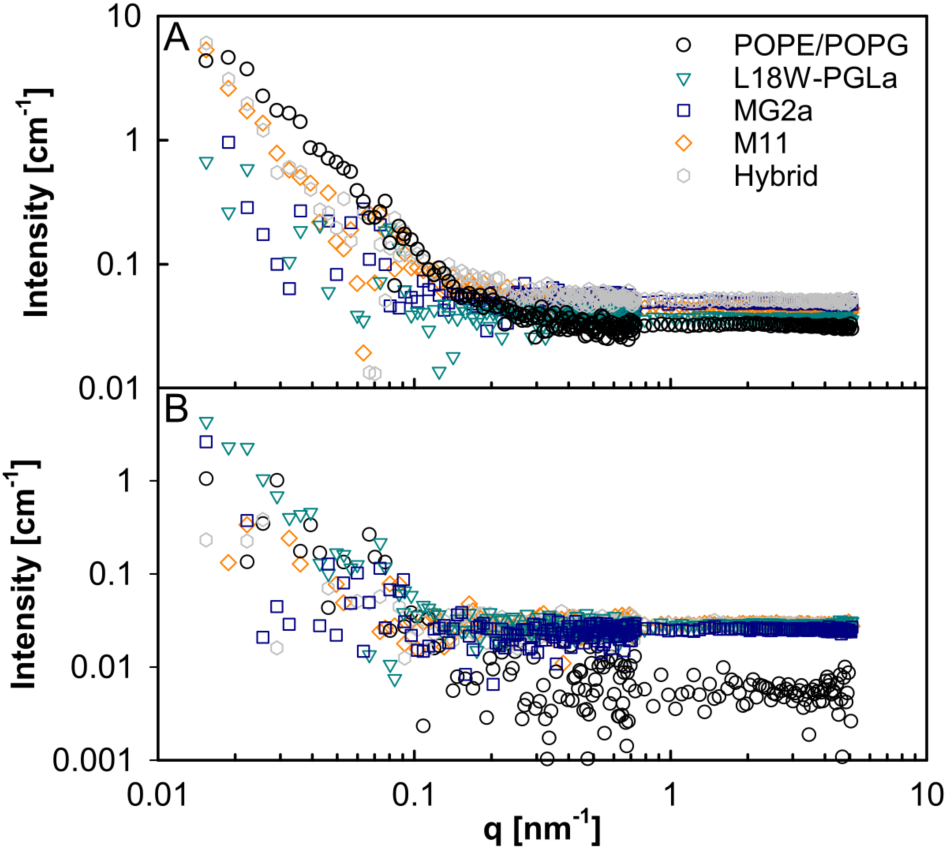
Zero-contrast SANS data of (POPEd31)/(POPG/POPGd31) (75)/(22/3) (Panel A) and the control sample (Panel B) in the absence (black circles) and presence of the peptides at 35°C and a P/L of 1:25.

#### Peptides transform LUVs to MLVs with collapsed layer spacing

In the following we focus on the end states (see the Methods section) of the systems well above the melting regime. SAXS data of the pure POPE/POPG samples showed a diffuse modulation of the scattered intensity, characteristic of positionally uncorrelated bilayers, as present in LUV dispersions which was analyzed in detail in the preceding paper (7). Equilibration of the LUVs with L18W-PGLa or MG2a (P/L = 1:50) did not affect the diffuse character of the scattering patterns (Fig. 3 *A*). In contrast, additional sharp peaks occurred for M11 and the peptide hybrid at the same P/L. The peaks can be ascribed to a single lamellar lattice of *d* = 49.8 Å (Fig. S2 *A*) in case of M11. The hybrid peptide induced a similar lamellar phase coexisting with a cubic Pn3m phase with a lattice constant of *a* = 100 Å and a prominent *q* _(1,1,0)_ ∼ 0.088 Å^-1^ peak (Fig. S2 *B*, Tab. S1). Note that we observed the Pn3m phase for the hybrid peptide already at P/L = 1 : 200 (Fig. 3 *B*, insert).

**Figure 3:**
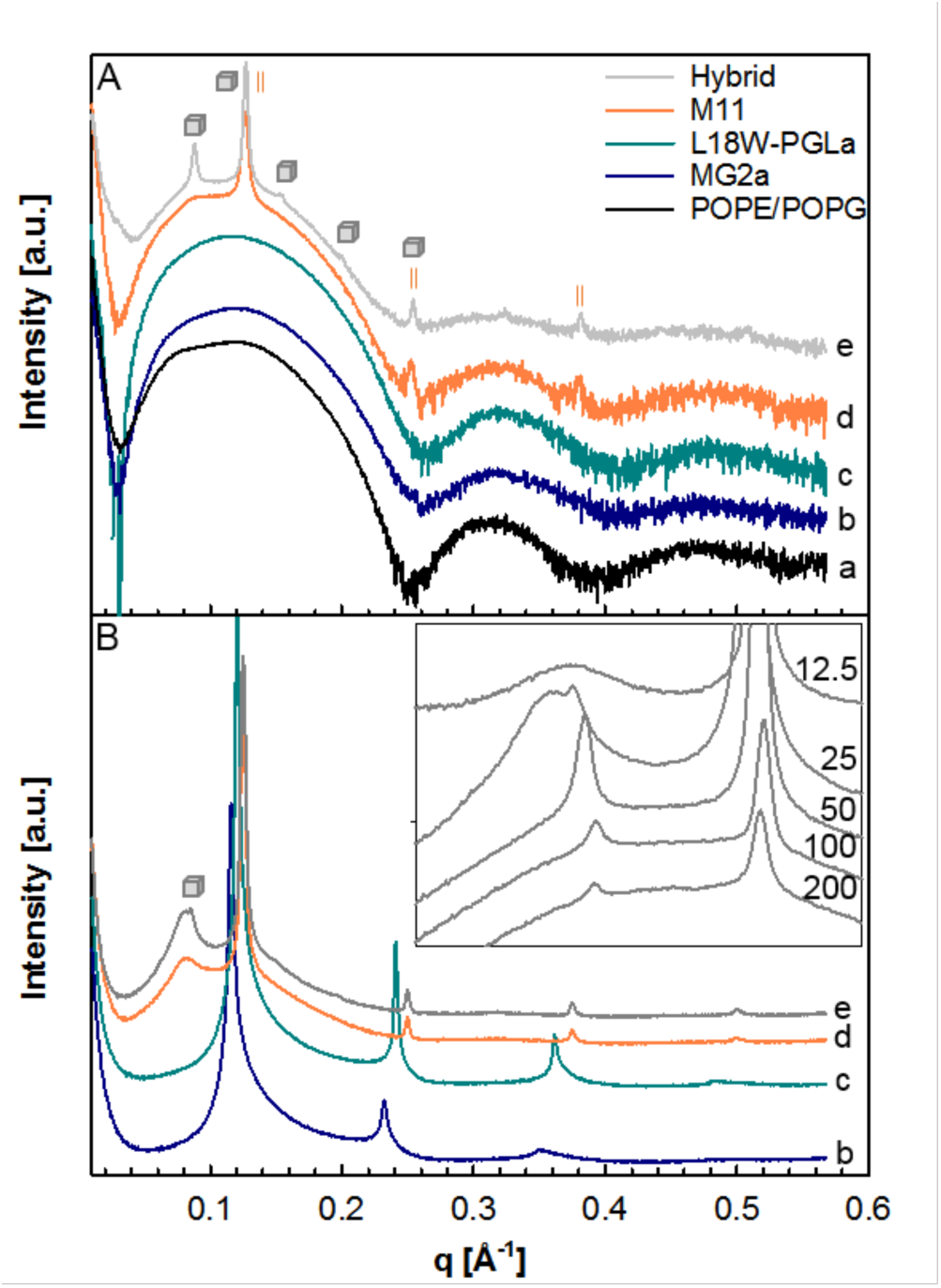
SAXS patterns of pure POPE/POPG LUVs (a) and in the presence of MG2a (b), L18W-PGLa (c), M11 (d) and the hybrid peptide (e) at P/L = 1:50 (panel A) and P/L = 1:25 (panel B), respectively at 35°C. The insert in panel B corresponds to a closeup of the scattering data in the low *q* region for the hybrid peptide. Numbers correspond to relative number of lipids for P/L, with P = 1.

Doubling the amount of peptide concentration (P/L = 1 : 25) also caused a formation of a multilamellar phase for the individual peptides (Fig. 3 *B*) with *D*^MG2a^ > *D*^L18W-PGLa^ > *D*^M11^ = *D*^hybrid^ (Tab. 1). While formation of a multilamellar aggregate was the only effect of L18W-PGLa or MG2a, their equimolar mixture induced an additional broad peak at *q* = 0.081 Å^-1^. The significant different shape of this peak as compared to the Bragg peaks originating from the lamellar phase clearly shows that it originates from different positional correlations. In fact, this peak was even present, though much weaker, at P/L = 1:50 at slightly higher *q*-values. Interestingly, we observed it also for the hybrid peptide at P/L = 1:25, where it partially overlapped with the (1,1,0)-reflection of the Pn3m phase (Fig. 3 *B*, insert). No signatures of a cubic phase were detected at P/L = 1:12.5, but the broad peak was clearly present.

The SAXS patterns observed at P/L = 1:25 for M11 and the hybrid peptide both displayed four lamellar diffraction orders (Fig. 3 *B*). This enabled us to calculate the steric bilayer thickness as detailed in the Methods section and the thickness of the water layer according to *D*_*W*_ = *D* – *D ′*_*B*_. Our results show very small *D*_*W*_ values of ∼ 7 Å (Tab. 1). That is, the formed lamellar aggregates are almost completely collapsed and most likely separated only by the steric size of partially membrane-inserted peptides. The lower amount of Bragg reflections observed in the presence of L18W-PGLa or MG2a individually impedes us from deriving electron density profiles with comparable resolution. However, the similarly small *D*-values suggest that also these lamellar aggregates are almost completely collapsed.

Increasing temperature leads to a linear decrease of the lamellar repeat distance with a similar rate for MG2a, L18W-PGLa and the hybrid peptide (Fig. 4), which is most likely due to a progressive thinning of the membrane, observed previously for other systems (see, e.g., (31)). The exception was M11, for which we observed the same *D* as for the hybrid peptide at low temperatures. However, *D* diverged from these data at 45°C and approached the *D* values of MG2a at 66°C. The difference between the hybrid and M11 is probably due to the decreased translational and configurational degrees of freedom for the hybrid, where both peptides are covalently coupled. In the case of M11 peptides interact only via weak forces, which seem to be overcome upon increasing temperature. Previously, we reported that MG2a inserts somewhat less deeply into the headgroup regime of POPE/POPG mixtures than L18W-PGLa (7). The temperature behavior of *D* for the equimolar mixture therefore indicates that the heterodimers dissociate for *T* > 45°C and peptides take up their ‘original’ positions within the bilayer.

**Figure 4:**
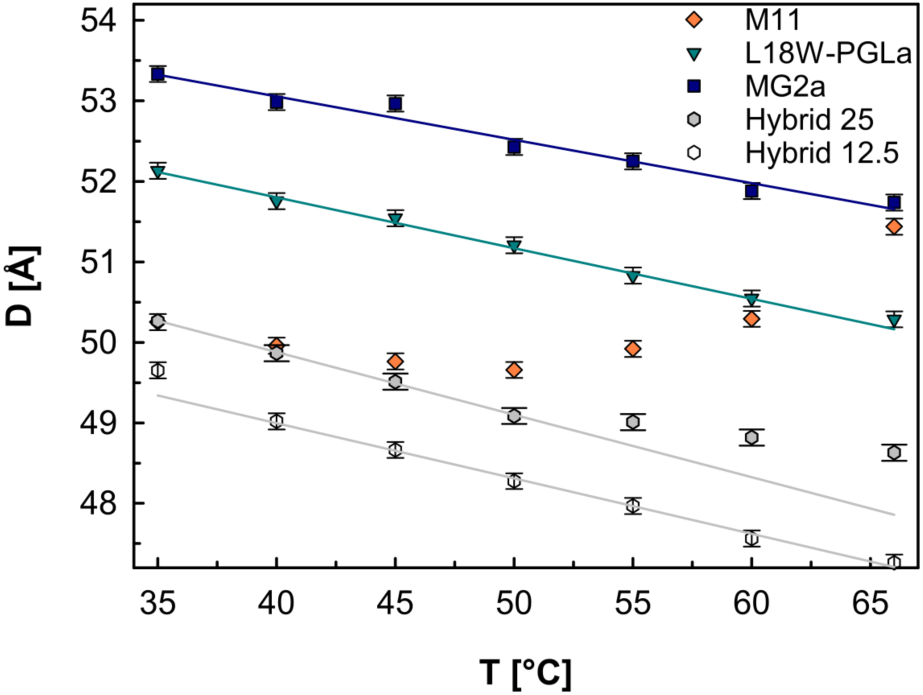
Temperature dependence of the lamellar repeat distance of POPE/POPG (3:1 mol/mol) multibilayers in the presence of magainins at P/L = 1:25 (full symbols) and P/L = 1:12.5 (open circle).

**Table 1:**
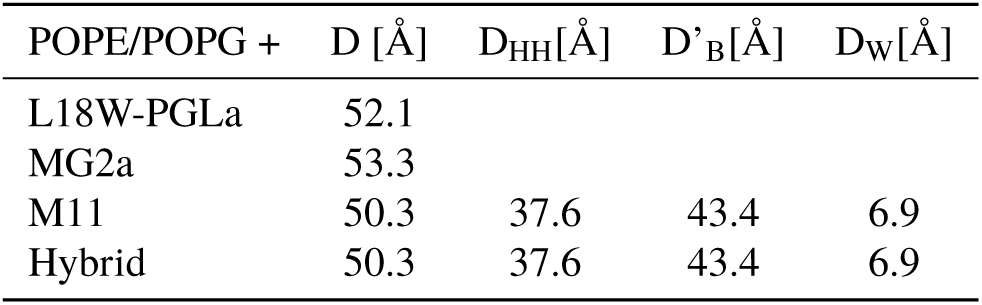
Structural parameters of peptide induced lamellar phases at P/L = 1:25.

It is not clear from SAXS experiments, whether the lamellar phases observed in the presence of peptides correspond to a stack of sheets or multilamellar vesicles (MLVs). To differentiate between these two types of aggregates we performed cryo-TEM. Adding either L18W-PGLa or MG2a (*P*/ *L* = 1 : 50) to POPE/POPG LUVs (Fig. 5 *A*) induced a significant variation of vesicle size without changing their lamellarity (Fig. 5 *B, C*). In contrast, the addition of M11 or the hybrid peptide induced clear signatures of MLVs (Fig. 5 *D, E*). We thus conclude that the peptides indeed lead to the formation of MLVs composed of tightly coupled bilayers. Note that the sharp peaks observed at P/L = 1:25 for the individual peptides (Fig. 3) indicate that MG2a and L18W-PGla are able to induce a LUV → MLV transition at sufficiently high concentration.

**Figure 5:**
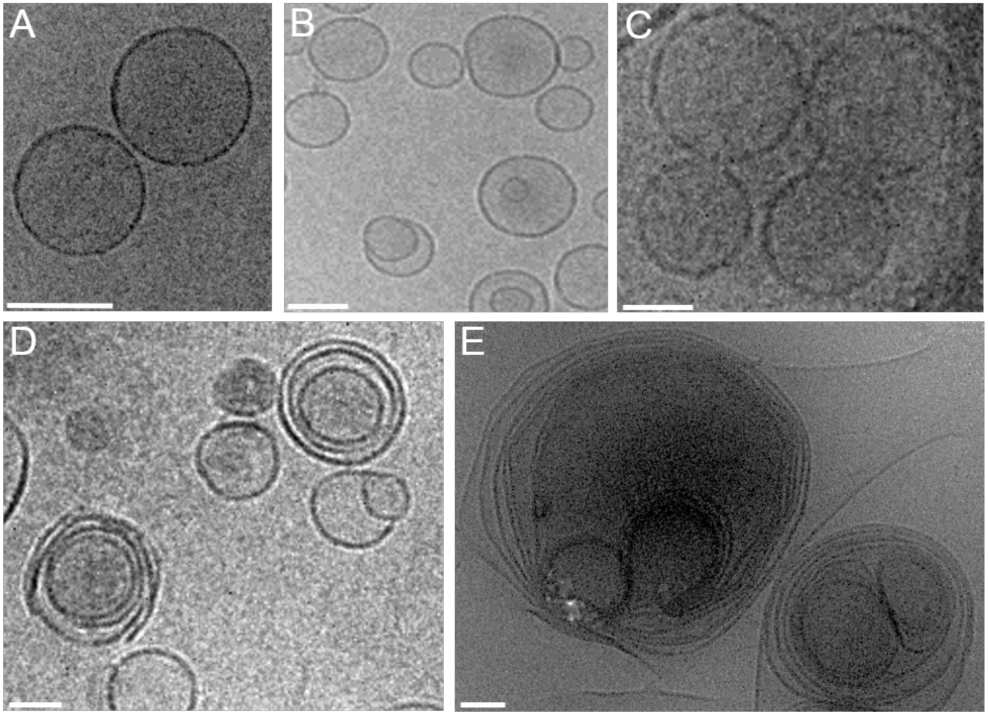
Cryo-TEM images of POPE/POPG (3:1 mol/mol) (A) and in the presence of the peptides (P/L = 1:50) L18W-PGLa (B), MG2a (C), M11 (D) and the hybrid peptide (E) at 35°C. All scale bars correspond to 100 nm.

### MD simulations

#### Peptide mixtures are able to form surface-bound tetramers in isolated bilayers

In the preceding paper, we found that MG2a and L18W-PGLa alone prefer to form surface-aligned monomers or antiparallel homodimers in POPE/POPG (3:1) bilayers (7). In contrast, equimolar mixtures of the peptides resulted in the preferential formation of parallel heterodimers in the same lipid bilayers and concentrations. Here we extended the coarse-grained simulation of the peptide mixture and found that the parallel heterodimers of MG2a and L18W-PGLa can further assemble into tetramers (Fig. 6). In these aggregates, all four peptides interact via their C-termini. Both MG2as are aligned completely parallel to the membrane surface, while L18W-PGLa peptides are slightly tilted with their C-termini reaching into the membrane hydrophobic core. Due to the amidation of the terminal carboxyl group and the lack of complementary charged species, the tetramer is held together mainly by hydrophobic interactions (see Fig. S7 for calculated interaction energies between individual residues).

**Figure 6:**
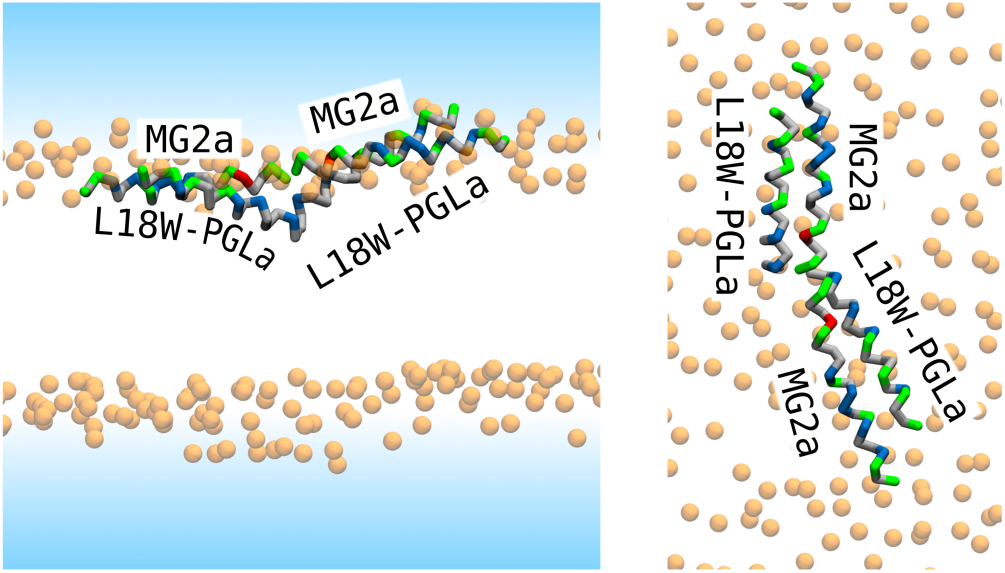
Representative simulation snapshots of two MG2a/L18W-PGLa parallel dimers forming a tetramer through the interaction of their C-termini. Side and top views on the tetramer are shown on left and right images, respectively. Lipid phosphate atoms are shown as orange spheres. Solvent is represented by a blue-shaded area and lipid tails are not shown for clarity. Peptide secondary structure is shown in cartoon representation and colored by residue type. Nonpolar: grey, polar: green, acidic: red, and basic: blue. Mag2a has a glutamic acid (red) in its sequence and is more polar, while L18W-PGLa is more hydrophobic.

Since previous NMR experiments do not provide information on the penetration depth of the two peptides (6) we also tested the possible formation of peptide aggregates within the membrane’s hydrophobic core. To do so, we inserted two parallel heterodimers in an alternating fashion in the membrane center, as shown in Fig. S8 *A, B*. Initially, the peptides were held in the position by restraints to equilibrate the membrane. Upon removal of these constraints, the peptide aggregate quickly fell apart and the peptides assumed surface-bound position (Fig. S8 (C)).

#### Formation of fibre-like structures in collapsed multibilayers

Motivated by our experimental evidence for the peptide-induced formation of MLVs with reduced intrinsic spacing, we prepared a stack of two membranes at P/L = 1:64 (see Methods section for details) and performed a 1 *µ*s long all-atom dynamics simulation, see Fig. S11 and 7. To mimic the experimental conditions, the bilayers centers were initially displaced by 50 Å as determined from our SAXS measurements. Due to the slow diffusion of peptides on membranes, the peptides were prepared as parallel dimers and placed randomly on the membrane. All dimers remained stable and interacted with lipids of the adjacent membrane. These peptides interactions lead to the formation of an alternating pattern of peptide nodes and solvent pockets (see Fig. 7). Any further aggregation of peptides was not observed within the 1 *µ*s time scale and all peptides remained within the initial membrane with their hydrophobic side facing the center of the bilayer. Note that the system size likely affected the size of the water pockets as water is constrained to remain within the simulation box.

**Figure 7:**
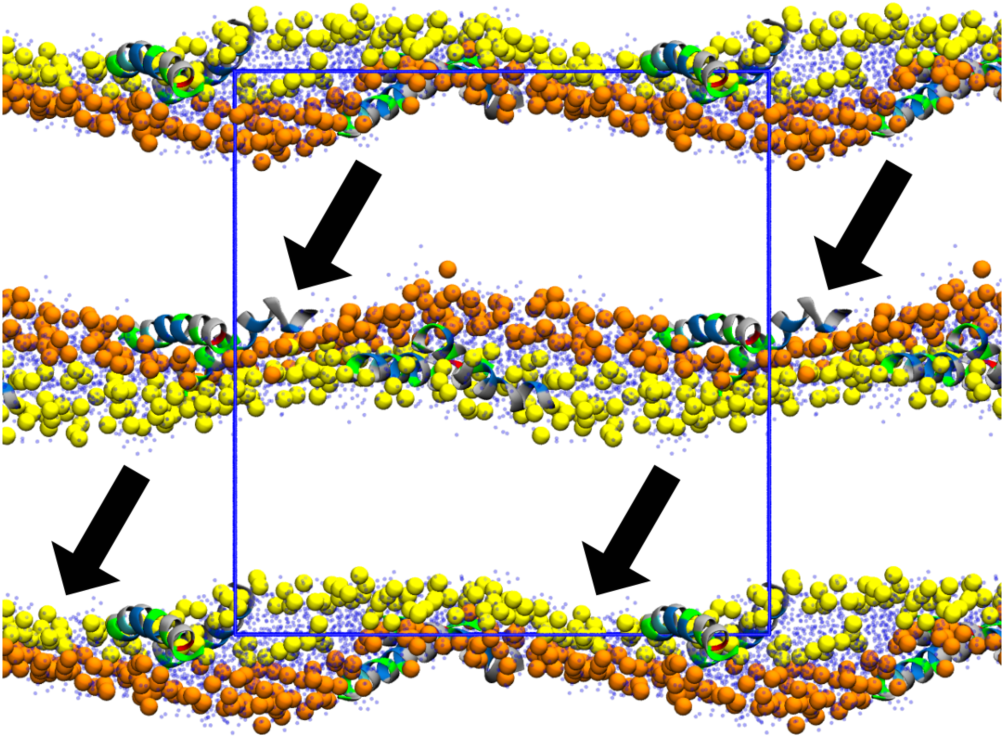
Final snapshot of two lipid bilayers after 1 *µ*s long all-atom simulation. Blue rectangle marks the simulation cell. Lipid phosphate atoms of first and second bilayers are shown as orange and yellow spheres, respectively. Lipid tails are not shown for clarity. The peptides’ secondary structures are displayed in cartoon representation and colored by residue type. Nonpolar: gray, polar: green, acidic: red, and basic: blue. Water oxygens are shown as semi-transparent dark-blue dots.

To verify that the observed behavior is not an artifact, we extended the lateral system size to 504-lipid bilayers and performed 20 *µ*s long coarse-grained Martini simulations in the presence and absence of L18W-PGLa and MG2a. No spontaneous membrane adhesion or appreciable interaction between the bilayers was observed for the pure membrane system (Fig. 8 *A*). Membrane adhesion with a concomitant formation of “pockets and nodes” was only observed in the presence of the peptide mixture. At P/L = 1:42, the peptides aggregated resulting in a height modulation of the membranes, where peptides were located at the nodes (Fig. 8 *B*), in agreement with all-atom simulations. The wavelength of the modulation is different from all-atom simulations and is likely to be affected by the system size. At higher peptide concentration (P/L=1:21), L18W-PGLa and MG2a formed fibril-like structures along the bilayer contact nodes (connected through the periodic boundary), as shown in Fig. 8 *C, D*. Interestingly, we also observed a temporary formation of membrane fusion stalk in the presence of the peptides (Fig. S12). The fusion stalk fell apart after the exchange of several lipids in tens of nanoseconds.

**Figure 8:**
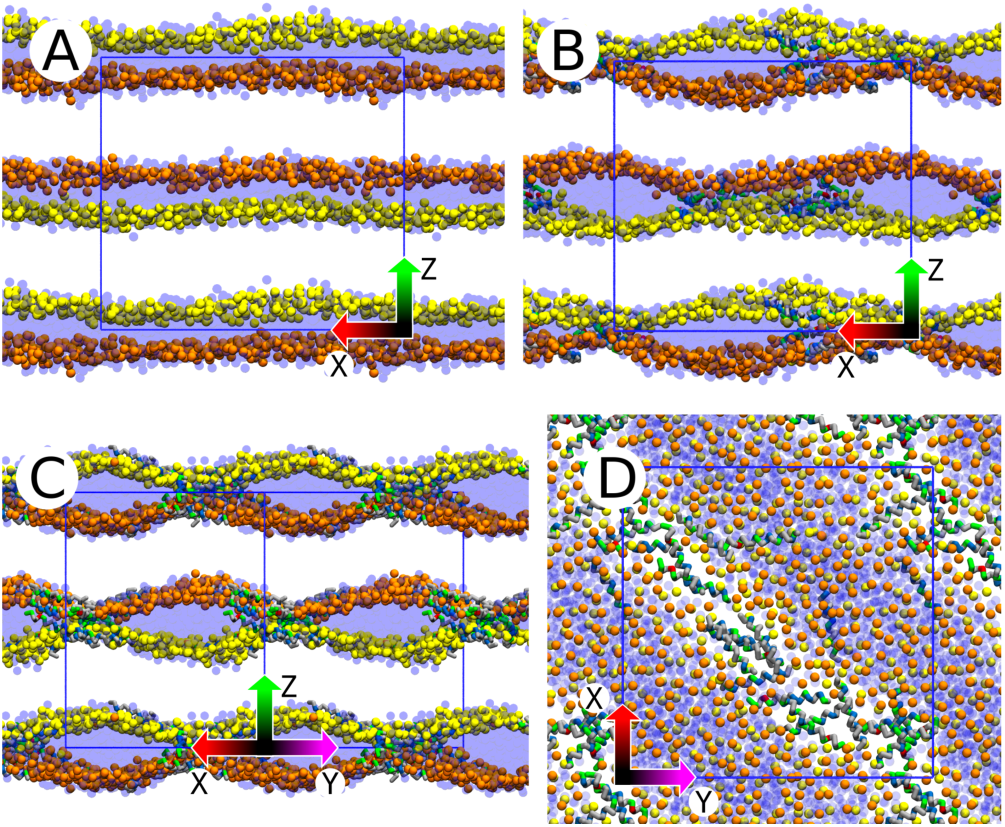
Last simulation snapshots of coarse-grained systems with two lipid bilayers and (A) without peptides, (B) with MG2a+L18W-PGLa at 1:42 peptide-to-lipid ratio, and (C, D) with MG2a+L18W-PGLa at 1:21 peptide-to-lipid ratio after 20 *µ*s. Arrows indicate the system axes: (X) red, (Y) magenta, and (Z) green. Blue rectangle marks the simulation cell: (A, B) view along the Y axis, (C) cell is rotated by 45° (because nodes formed in that direction this time), and (D) top view along the Z axis. Lipid phosphate atoms of the first and second bilayers are shown as orange and yellow spheres, respectively. Lipid tails are not displayed for clarity. Peptide secondary structure is shown in cartoon representation and colored by residue type. Nonpolar: gray, polar: green, acidic: red, and basic: blue. Water beads are shown as semi-transparent dark-blue spheres.

To investigate the effect of periodic boundary conditions, we prepared a much larger system with a stack of membranes each composed of 2016 lipid molecules. The peptides were added at random initial positions and orientations (P/L = 1 : 21), but distributed equally on the membrane surfaces. Fig. S9 *A* shows the initial configuration from the production dynamics simulation. During the simulation, a fusion stalk formed between two bilayers and remained stable for the remaining duration of the simulation, i.e., 100 *µ*s. Fig. S9B shows the snapshot from the end of the simulation. Note that even during the stalk formation the peptides remained parallel or slightly tilted to the membrane plane. Irrespective of the fusion stalk, which occurred only at a single location of the simulated membrane stack, the peptides aggregated and formed fibril-like structures, analogously to the 504-lipid simulation (Fig. S10).

In order to interrogate, whether the presence of peptides in both leaflets of opposing bilayers is a necessary condition for bilayer adhesion we performed a coarse-grained simulation of a similar-sized system but placing the peptides in one leaflet of each bilayer, only. We again observed a membrane stalk. Interestingly, its formation was accompanied by a translocation of peptides between the membranes. Specifically, stalk-formation was initiated from a heterodimer by moving a MG2a toward the interbilayer space (Fig. 9). This caused one of the lipids from the opposite membrane to insert its tail between the peptide dimer. The insertion was stabilized by the hydrophobic interactions of the hydrocarbon chain with the phenylalanine residues of MG2a at position 12 and 16. Subsequently, the heterodimer split at the N-terminus leading to the formation of a fusion stalk, which was further stabilized by other peptides, which diffused into the stalk. Similar process of stalk formation was observed it in three independent simulations.

**Figure 9:**
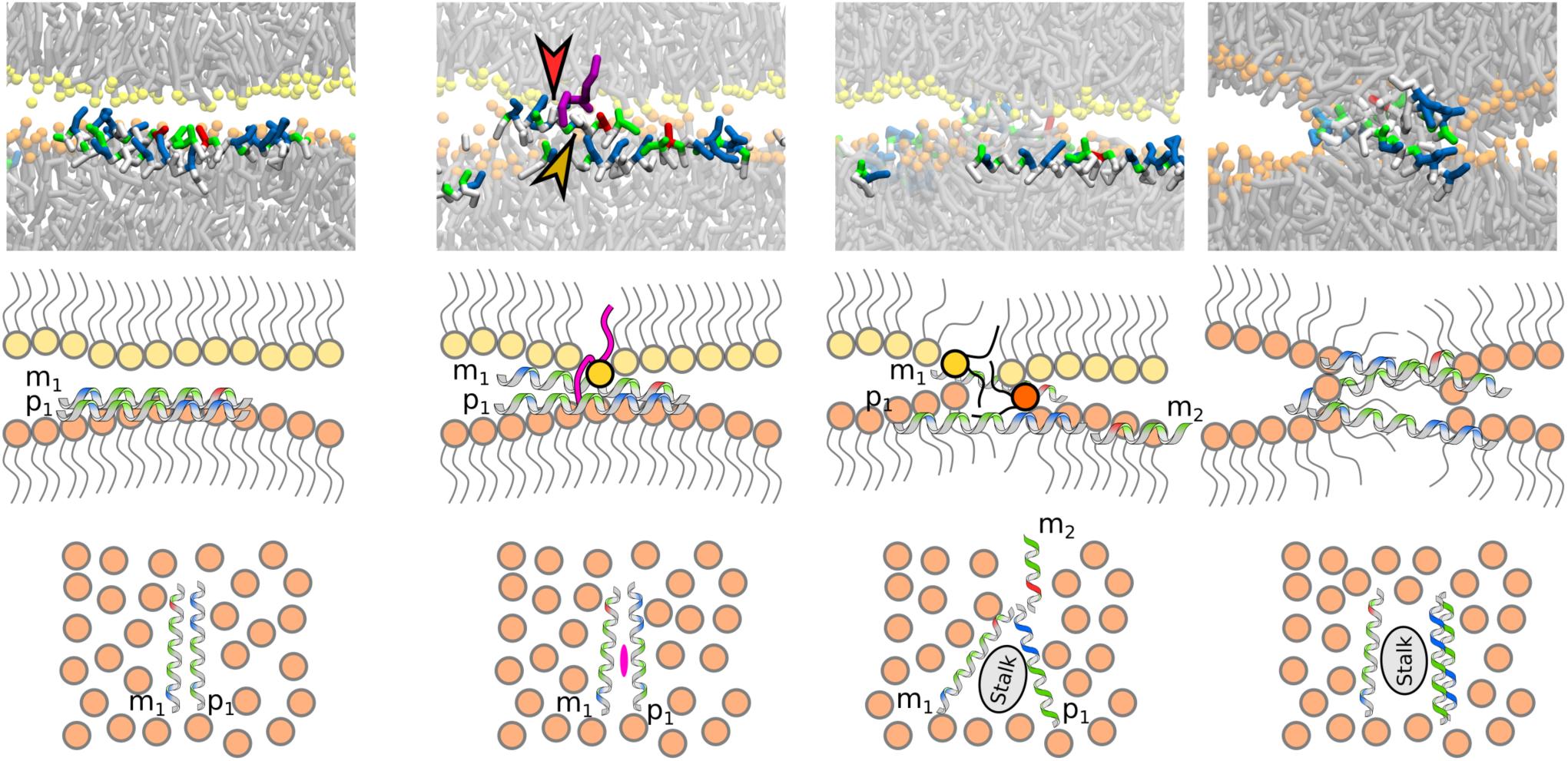
Simulation snapshots (top) and corresponding illustrative images from the side (middle) and top view (bottom) capturing the process of bilayer fusion. 1) MG2a (m1) and L18W-PGLa (p1) peptides arranged in a parallel dimer. 2) The bilayer fusion was initiated by one lipid (highlighted in purple). Single lipid tail was interacting with Phe12 (red arrow) and Phe16 (dark-yellow arrow) of the MG2a peptide and connected both bilayers. 3) MG2a and L18W-PGLa peptides assumed a V-shaped configuration that caused the formation of a fusion stalk. 4) Several peptides crossed from one bilayer to another and lipids from the two bilayers started to mix through the stalk. Later in the simulation, two L18W-PGLa peptides formed an antiparallel homodimer and the fusion stalk further expanded. One MG2a peptide was stabilizing the stalk from the other side. The stalk remained stable until the end of the 20 *µ*s long simulation. In first three columns, phosphate beads of the first and second bilayer are shown in yellow and orange, respectively. In the most right column, all phosphate beads are shown orange due to lipid mixing.

## DISCUSSION

We used a combination of experimental and computational techniques to demonstrate that equimolar mixtures of L18W-PGLa and MG2a induce intricate topological changes in membranes composed of POPE/POPG (3:1 mol/mol) at synergistic peptide/lipid ratios. The peptide mixture caused a transformation of LUVs into MLVs with a collapsed interbilayer distance as evidenced by SAXS and cryo-TEM (Figs. 3, 5). MG2a and L18W-PGLa individually were also found to induce this transformation, however, only at doubled concentration. At the molecular level, our coarse-grained MD simulations showed that the formation of parallel L18W-PGLa/MG2a heterodimers observed in a previous report (7) progresses to a tetramer formation (or double heterodimers) stabilized by C-termini interactions, which can further assemble into fibril-like structures in presence of two close membranes (Figs. 6 - 8). These fibril-like structures are aligned parallel to the membrane surface and are sandwiched between the collapsed multibilayers, i.e. the peptides couple adjacent bilayers by peptide-peptide and peptide-lipid interactions. Interestingly, electrostatic interaction between POPG and the cationic residues of MG2a and L18W-PGLa does not induce lipid domain formation as evidenced by zero-contrast SANS (Fig. 2). Instead, the splitting of the main phase transition of the aggregates into multiple melting transitions appears to be due to a non-homogeneous distribution of peptides.(Fig. 1). We note, however, that the overall melting process was significantly less cooperative for the peptide mixture and the hybrid peptide, i.e., in the presence of their aggregates.

The formation of fibril-like structures induced a curvature modulation of membranes with peptides forming the connections between the membranes. These peptides aggregates were able to initiate the formation of fusion stalks (Figs. 9, S12). The smallest peptide aggregate, for which we observed stalk formation, was a parallel heterodimer, inserted only in one membrane leaflet, which induced a fusion to the proximal peptide-free bilayer. Interestingly, peptides remained parallel or slightly tilted from the membrane plane even during the stalk formation.

Stalks are well-known precursors to membrane fusion (32–34) and cubic phase formation (32). Indeed, we observed a Pn3m cubic phase coexisting with the collapsed multibilayers in case of chemically-linked heterdimers by SAXS (Fig. 3), thus supporting MD results. At elevated concentration of the hybrid peptide (P/L = 1:25), we observed an additional broad peak indicating the presence of an additional phase coexisting with the Pn3m and lamellar phases. The Pn3m completely vanished at P/L = 1:12.5, and the scattering pattern exhibited just a broad peak as well as the peaks arising from the collapsed multibilayers. A similar broad Bragg reflection was also observed for L18W-PGLa/MG2a mixtures, but not for the individual peptides (Fig. 3).

There are two possible scenarios for the origin of this broad peak, the first one being the peptide-induced curvature modulation of the lamellar phase. In this case, the peak would emanate from the wavelength of the modulation. Somewhat similar patterns have been reported for sandwiched structures formed by DNA/surfactant complexes under specific conditions (35), where DNA aligned between the bilayers. A second scenario explaining the presence of the broad peak is the formation of a sponge phase. Sponge phases can be viewed as molten bicontinuous cubic phases, with a characteristic low-angle peak, whose position relates to the average distance between the water channels (36–38). Likewise, the *D*_(1,1,0)_ of the Pn3m phase, gives the distance between the pores. The facts that (i) the *D*-spacing of the broad peak is larger than *D*_(1,1,0)_ in the presence of the hybrid peptide at P/L = 1:25 and (ii) the Pn3m phase vanishes upon further increasing the peptide concentration (Fig. 3 *B*, insert) indeed support the formation of a sponge phase, which analogously to Pn3m consists of a network of water channels, but lacks 3D crystalline-like positional correlations. In other words, increasing hybrid peptide concentration appears to lead to an overall decrease of membrane curvature causing a melting of the Pn3m phase into a spongy structure. The similar position and shape of the broad Bragg peak formed in presence of M11 suggest also the formation of a sponge phase in the synergistic regime of the peptide mixture. Previously, Silva et al. reported a LUV → collapsed MLV transition for the same lipid mixture in the presence of cecropin A (39). However, no coexisting sponge phase or other curvature-related structures were observed. We found black featureless objects in samples with the equimolar mixture or the hybrid peptide (Fig. S13). These structures could contain cubic or sponge structures, but are too large/thick to be analyzed by cryo-TEM.

Finally, it is important to align our present findings with our previously reported leakage data for the same lipid system (4). Both L18W-PGLa and MG2a caused only minor release of fluorescent dye, whereas their equimolar mixture and the hybrid peptide induced significant leakage. Our present experiments revealed the formation of multibilayers with a collapsed *D*-spacing at high peptide concentration for all peptides, including their equimolar mixture (Figs. 3). Therefore, a LUV→ MLV transition does not seem to be a sufficient condition for enhanced dye leakage. This suggests that there is a pathway transforming LUVs to MLVs without leaking their content. In contrast, synergistically increased dye release is coupled to increased membrane curvature in the presence of fibril-like peptide structures leading to fusion events and the development of sponge or cubic phases. Importantly, fusion events could be leaky (40, 41) and both above phases are highly leaky structures able to account for the dye release. Additionally, the significant difference between *D*_(1,1,0)_ ∼ 73 Å of the Pn3m phase and the *D* ∼ 50 Å of the lamellar phase (Tab. 1) shows that there is no epitaxial relationship between the cubic and the lamellar phases. This indicates the formation of a defect-rich zone bridging the two structures, which could also give rise to significant membrane leakage (see, e.g., (38) for a detailed discussion).

An important aspect of our results is that both peptides remained in a surface-aligned topology even during the formation of fusion-stalks. This is specific to the significant negative intrinsic curvature of POPE, which leads to a tight packing of the bilayer’s polar/apolar interface and a high energy barrier for peptide translocation (4). Therefore other membrane mimics, lacking contributions of lipids with negative intrinsic curvature – e.g. if lipids not occurring in bacteria are used –, may allow for a full insertion of peptides and the formation of specific transmembrane pores as suggested previously for MG2a and PGLa (1, 2). Pores of sponge or cubic phases essentially result from bridging two close bilayers and are consequently distinct from peptide transmembrane pores. Our results agree in part with the suggested mesophase-like peptide arrangement (3), which in a way are similar to the fibril-like structures observed here, although effects on membrane curvature were not considered. However, effective discharge of dyes from LUVs correlates with the formation of sponge or cubic phases.

Correlating our present findings to the biological activity of the peptides (4) is not straightforward, due significant differences between Gram-negative bacteria and here studied lipid vesicles. We are not aware of any observation of peptide-induced non-lamellar membrane structures in bacteria. However, inner mitochondrial membranes have been reported to form cubic phases under stress conditions (42). Moreover, bacterial lipid extracts have been shown to form cubic phase in the presence of surface-inserted AMPs (43). Observations of cubic phases formed by membranes enriched with lipids with negative intrinsic curvature (such as phosphatidylethanolamine and cardiolipin) are in line with our findings. Aggregates, as observed for L18W-PGLa and MG2a, seem to facilitate this effect leading to sponge or cubic phases even in the absence of cardiolipin, which might be even be exploited by PGLa and MG2a in bacteria. Moreover, constantly ongoing membrane remodelling processes such as bacterial endocytosis (44) or cell division provide membrane contact points for peptides to intercept these processes possibly by inducing non-lamellar membrane structures.

## CONCLUSION

The synergistic mechanism of MG2a and L18W-PGLa is a complex process which evolves over several length scales. MG2a and L18W-PGla are able to dimerize already at low peptide concentrations (7), which progresses upon increasing their content into fibril-like structures inducing cross-links between proximate bilayers leading to the occurrence of fusion stalks. The membrane with synergistic peptides evolve over time into a sponge phase correlating with synergistic leakage of the vesicles as observed previously (4). If the peptides are strongly coupled, as in the case of the MG2a-L18W-PGLa hybrid, a cubic phase with high membrane curvature forms already at peptide concentrations as low as P/L = 1:200 (Fig. 3 *B*), tying in with the hybrid’s even higher activity (see Fig. 4 *A* in (4)). One of remaining issue to address regarding the synergistic mechanism of PGLa and MG2a is the sequence of events, i.e. the kinetics of their activity. We are currently performing such studies in our laboratories.

## Supporting information

Supportin Information

## AUTHOR CONTRIBUTIONS

I.K. carried out all simulations, analyzed the data, and wrote the article. M.P. performed SAXS, SANS, DSC experiments, analyzed the data, and wrote the article. S.P. performed SANS experiments and I.L-P. performed and analyzed TEM experiments. R.V., K.L. and G.P. designed the research and wrote the article.

## ACKNOWLEDGMENTS

We thank Frederick A. Heberle for assistance in designing the zero-contrast SANS experiments and Michael Rappolt for helpful insights regarding cubic and sponge phases. This work was supported by the Czech Science Foundation (grant 17-11571S to R.V.), the Austrian Science Funds FWF (project No. I1763-B21 to K.L.), Grant Agency of Masaryk University (MUNI/G/ 1100/2016), and the CEITEC 2020 (LQ1601) project with financial contribution made by the Ministry of Education, Youths and Sports of the Czech Republic within special support paid from the National Programme for Sustainability II funds. Computational resources were provided by the CESNET LM2015042 and the CERIT Scientific Cloud LM2015085, provided under the programme ‘Projects of Large Research, Development, and Innovations Infrastructures’. This work was supported by The Ministry of Education, Youth and Sports from the Large Infrastructures for Research, Experimental Development and Innovations project ‘IT4Innovations National Supercomputing Center – LM2015070’. We acknowledge SOLEIL for provision of synchrotron radiation facilities and we would like to thank Javier Perez for assistance in using beamline SWING.

