## Supplementary material for "Magainin 2 and PGLa in Bacterial Membrane Mimics II: Membrane Fusion and Sponge Phase Formation": Supportin Information

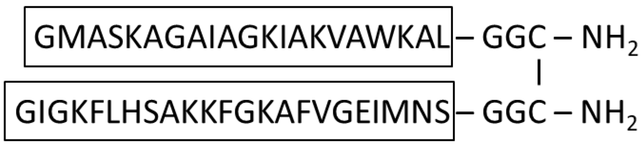


Figure S1: Chemical structure of the L18W-PGLa–MG2a hybrid peptide made of covalently bound peptides L18W-PGLa and MG2a.

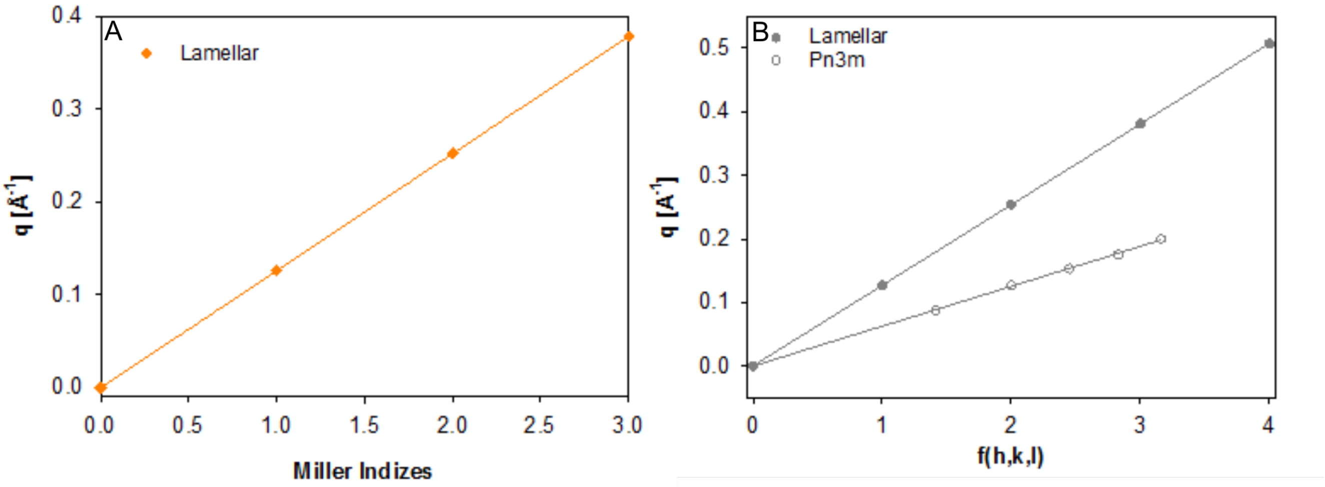


Figure S2: Indexation plots for the lamellar phase formed in the presence of M11 (A) and the coexisting lamellar and cubic phases formed by the chemically linked hybrid peptide (B) at a P/L of 1:50.

Table S1: Observed reflections of the lamellar and Pn3m phases.

| lamellar (h) | Pn3m (h,k,l) |
|---|---|
| 1 | 1,1,0 |
| 2 | 2,0,0 |
| 3 | 2,1,1 |
| 4 | 2,2,0 |
| | 3,1,0 |

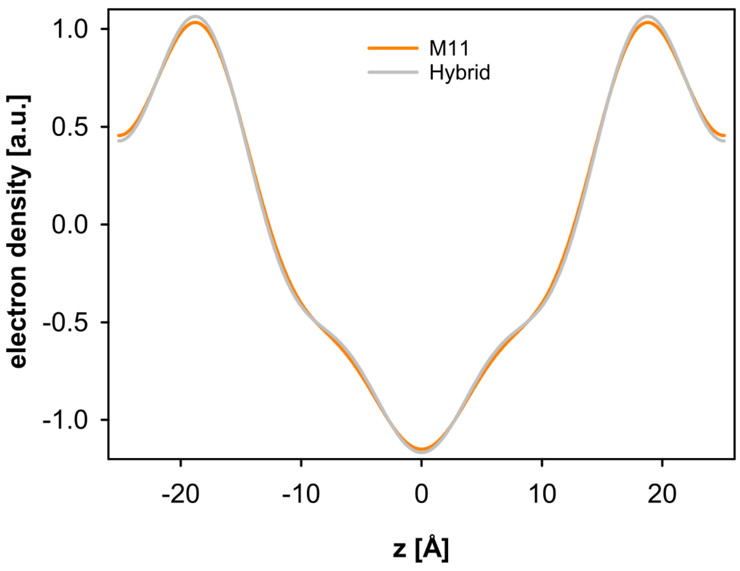


Figure S3: Electron density of 100 nm unilamellar vesicles composed of POPE/POPG (3:1 mol/mol) in the presence of M11 and the hybrid peptide at a P/L of 1/25 and 35°C.

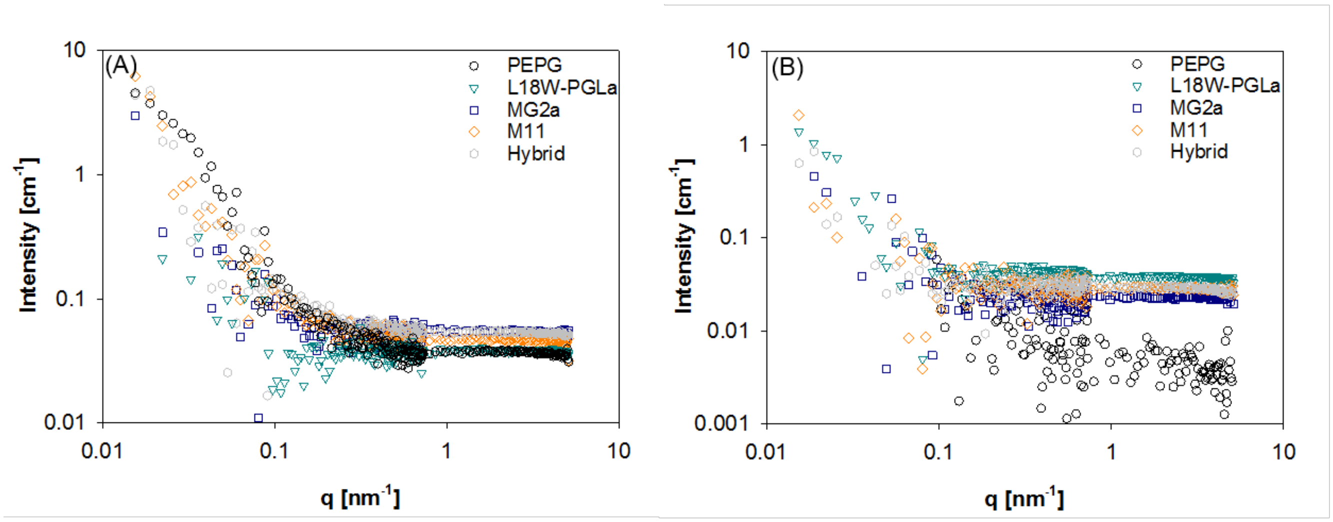


Figure S4: Zero-contrast SANS data of (POPEd31)/(POPG/POPGd31) (75)/(22/3) (Panel A) and the control sample (Panel B) in the absence (black circles) and presence of the peptides at 20°C (A) and 22°C (B) at a P/L of 1:25.

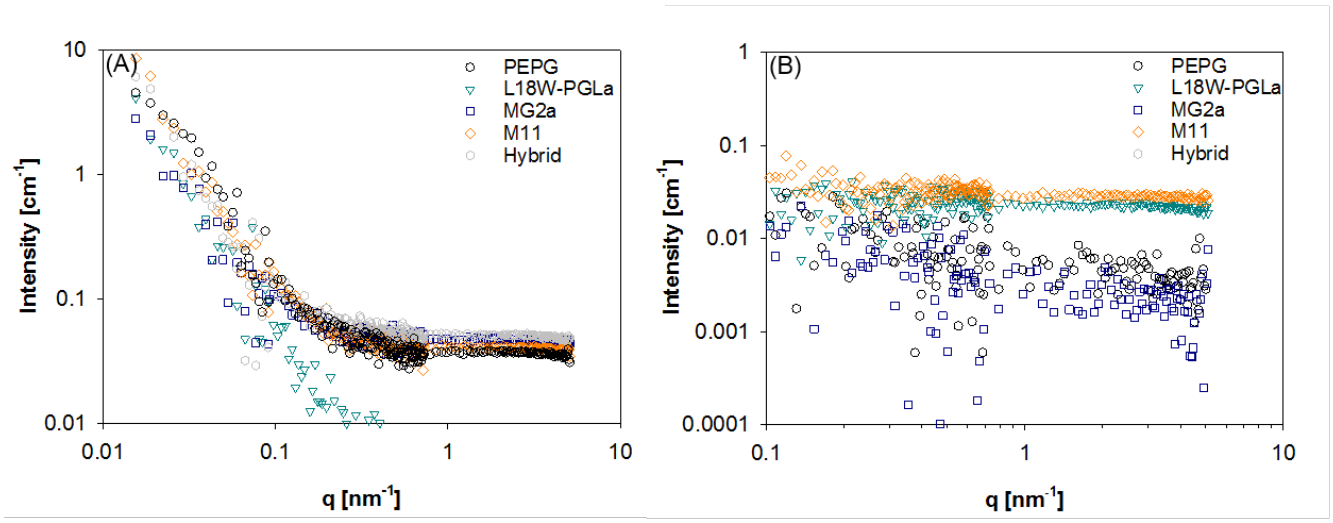


Figure S5: Zero-contrast SANS data of (POPEd31)/(POPG/POPGd31) (75)/(22/3) (Panel A) and the control sample (Panel B) in the absence (black circles) and presence of the peptides at 20°C (A) and 22°C (B) at a P/L of 1:50.

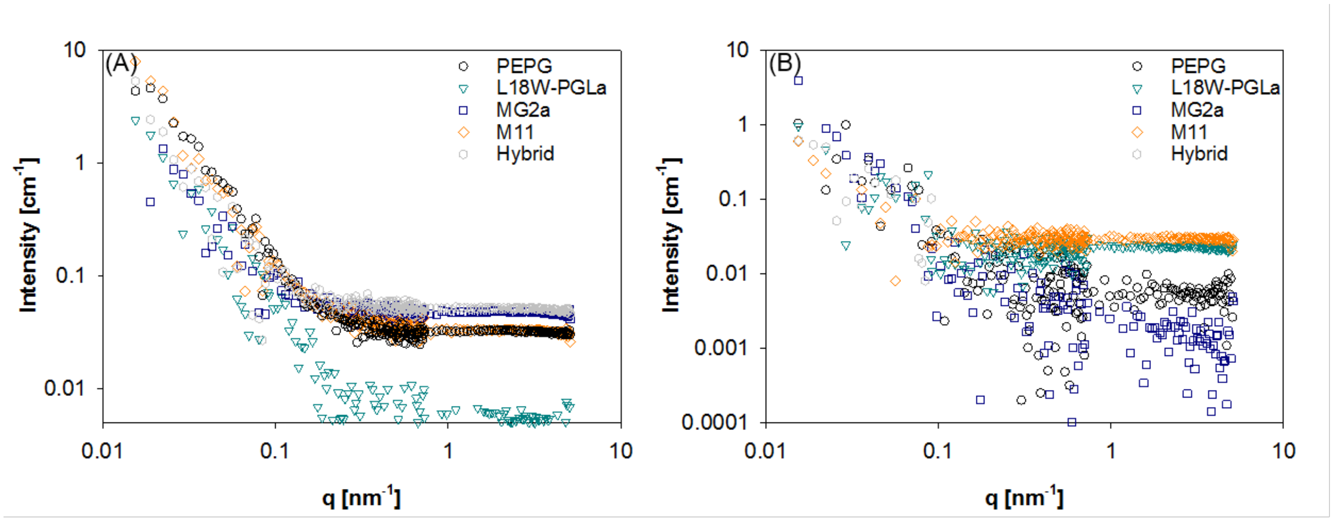


Figure S6: Zero-contrast SANS data of (POPEd31)/(POPG/POPGd31) (75)/(22/3) (Panel A) and the control sample (Panel B) in the absence (black circles) and presence of the peptides at 35°C° at a P/L of 1:50.

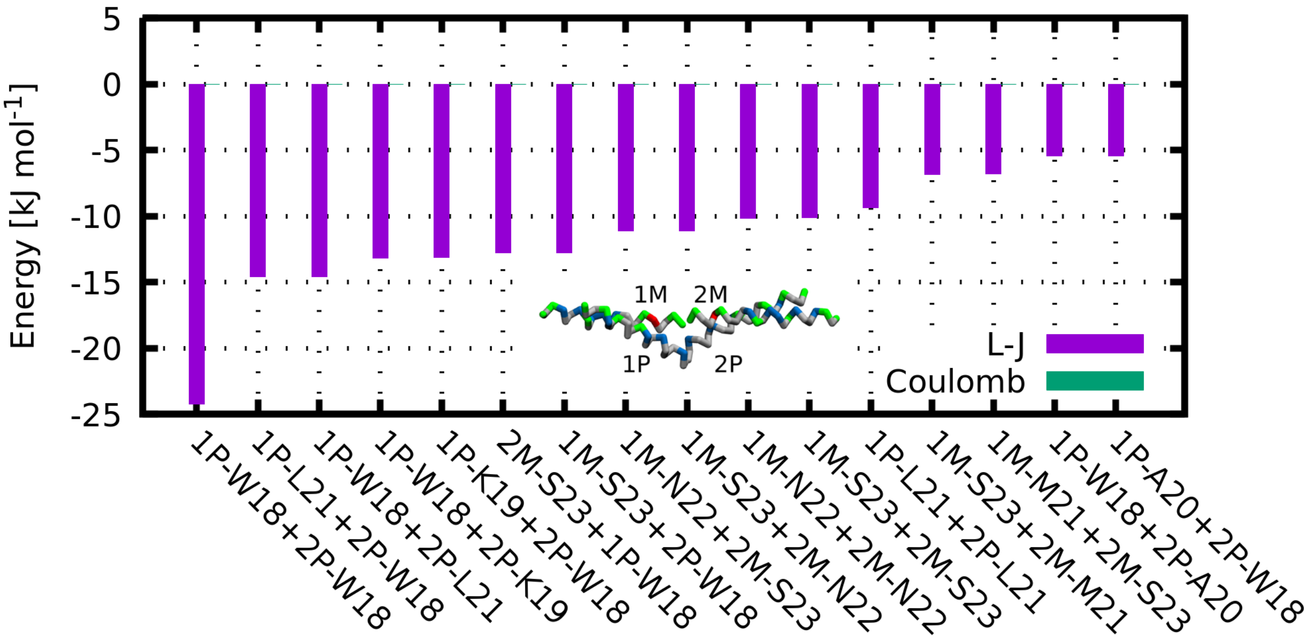


Figure S7: Calculated interaction energies between residues in opposing L18W-PGLa/MG2a heterodimers. Using single-letter amino acid codes, only residues with interaction energies larger than 5 kJ mol$^{-1}$ are shown. MG2a and L18W-PGLaW are abbreviated as ’M’ and ’P’, respectively. The inset shows a representative structure of the tetramer formed by the heterodimers pointing with C-termini in the center. Purple and green bars represent Lennard-Jones and Coulomb interactions, respectively.

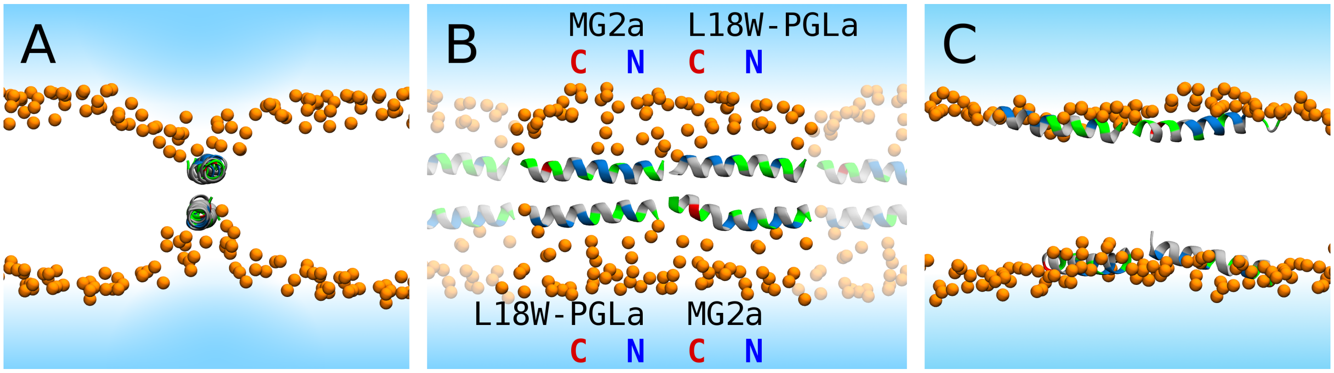


Figure S8: Initial and final simulation snapshots of a peptide fiber in the middle of the membrane. Initial system configuration with view along the fiber axis (A) and a sideview – rotated by 90° – (B). Periodic images are semi-transparent. (C) Final snapshot after 330 ns of production dynamics. Lipid phosphate atoms are shown as orange spheres. Lipid tails are not shown for clarity. The peptide secondary structure is shown in cartoon representation and colored by residue type. Nonpolar: gray, polar: green, acidic: red, and basic: blue.

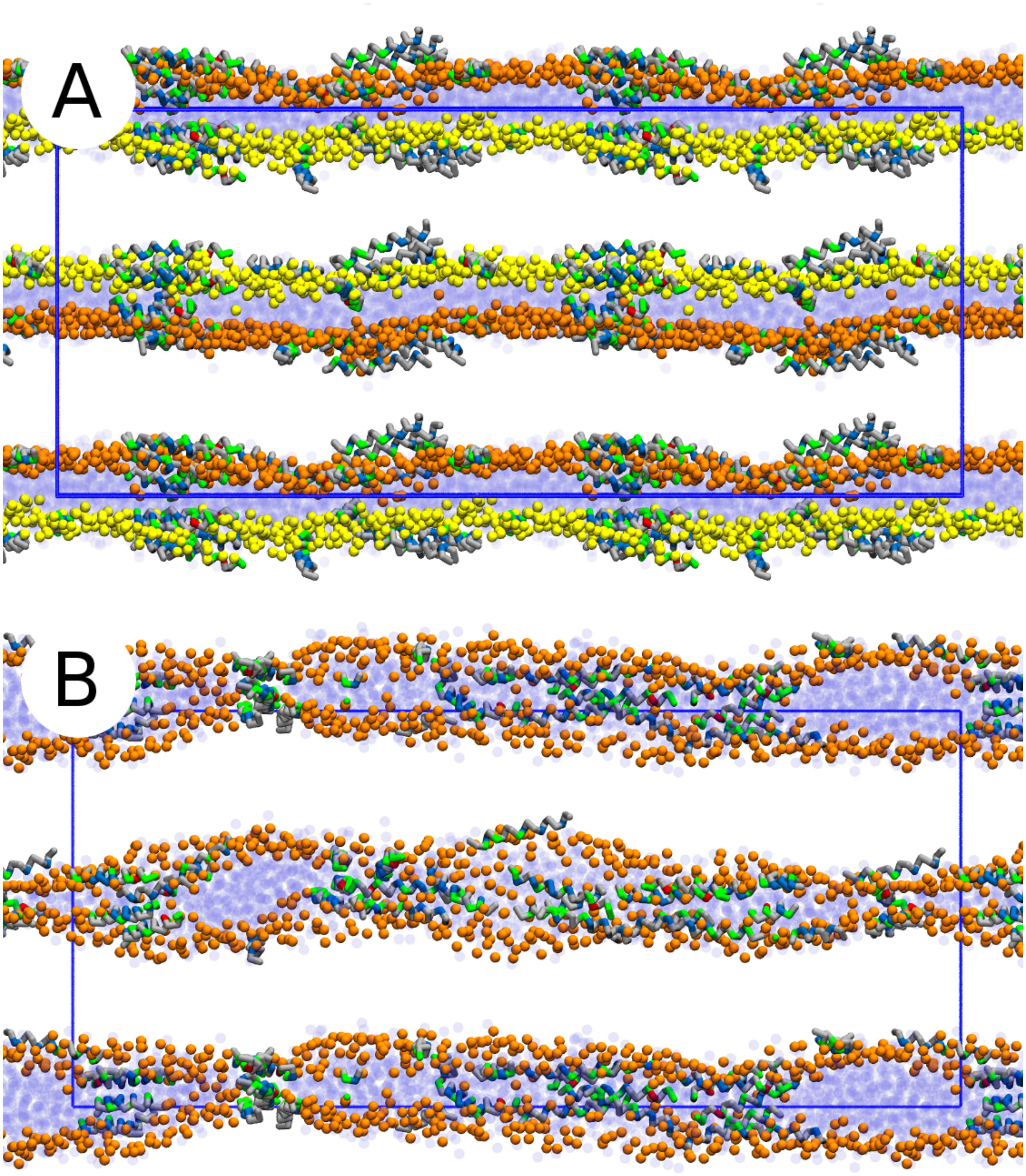


Figure S9: (A) Initial snapshot from the production simulation of a stack of two lipid bilayers with adsorbed peptides at 1:21 P/L. (B) Cut through the simulated system from the final simulation configuration after 100 $\mu$s. Due to the formation of a stable membrane stalk, lipids were mixed between bilayers. Lipid phosphate atoms of first and second bilayers before mixing are shown as orange and yellow spheres, respectively. Lipid tails are not shown for clarity. Nonpolar: gray, polar: green, acidic: red, and basic: blue. Water beads are shown as semi-transparent dark-blue spheres.

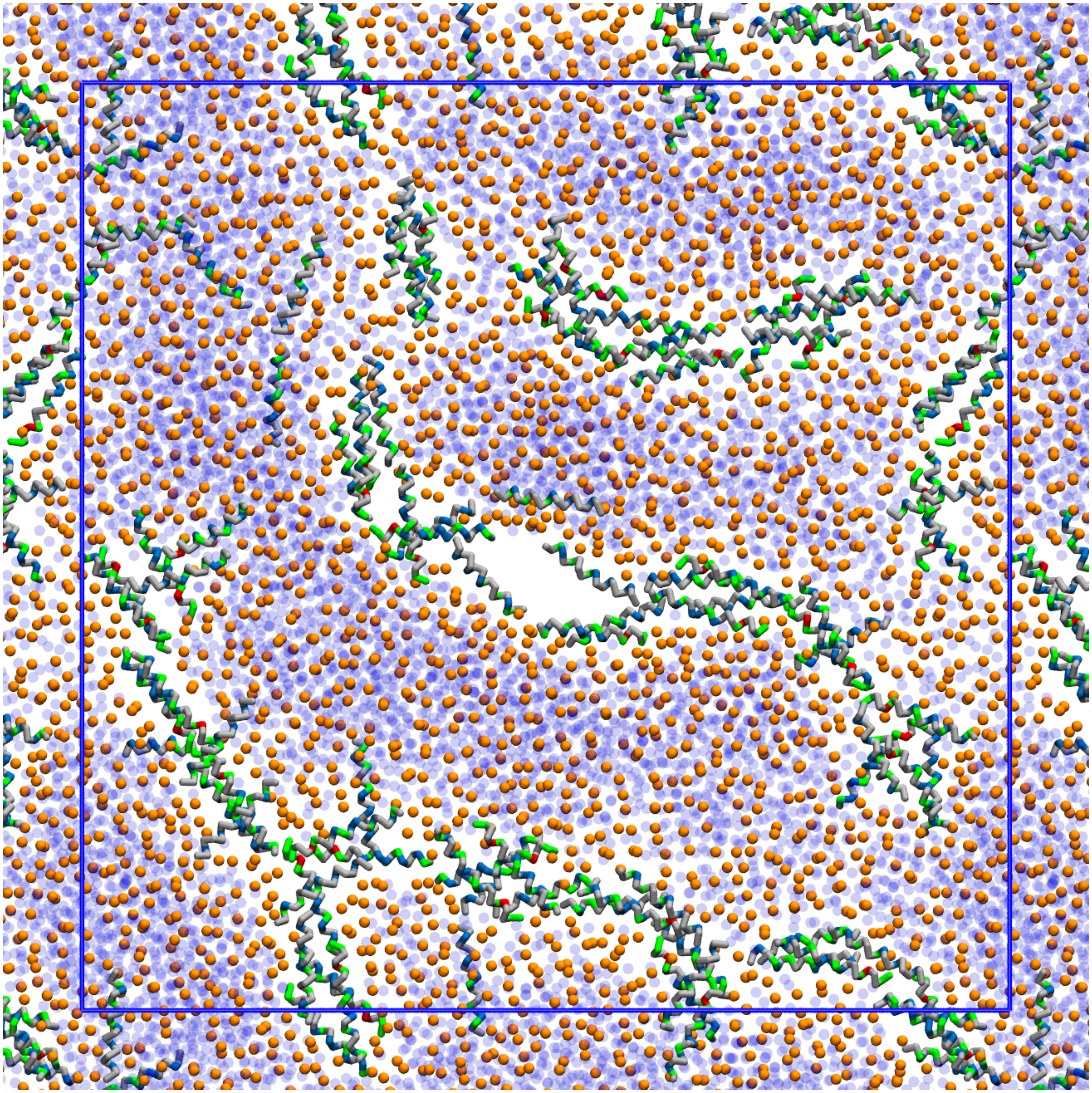

Figure S10: Top view on the interface between two lipid bilayers with adsorbed peptides at P/L= 1:21. The snapshot was taken from the end of a 100 $\mu$s. Peptides formed fiber-like structures and brought the membrane leaflets closer together. A stable fusion stalk is in the center of the image, surrounded by the peptides. Lipid phosphate atoms are shown as orange spheres. Lipid tails are not shown for clarity. Nonpolar: gray, polar: green, acidic: red, and basic: blue. Water beads are shown as semi-transparent dark-blue spheres.

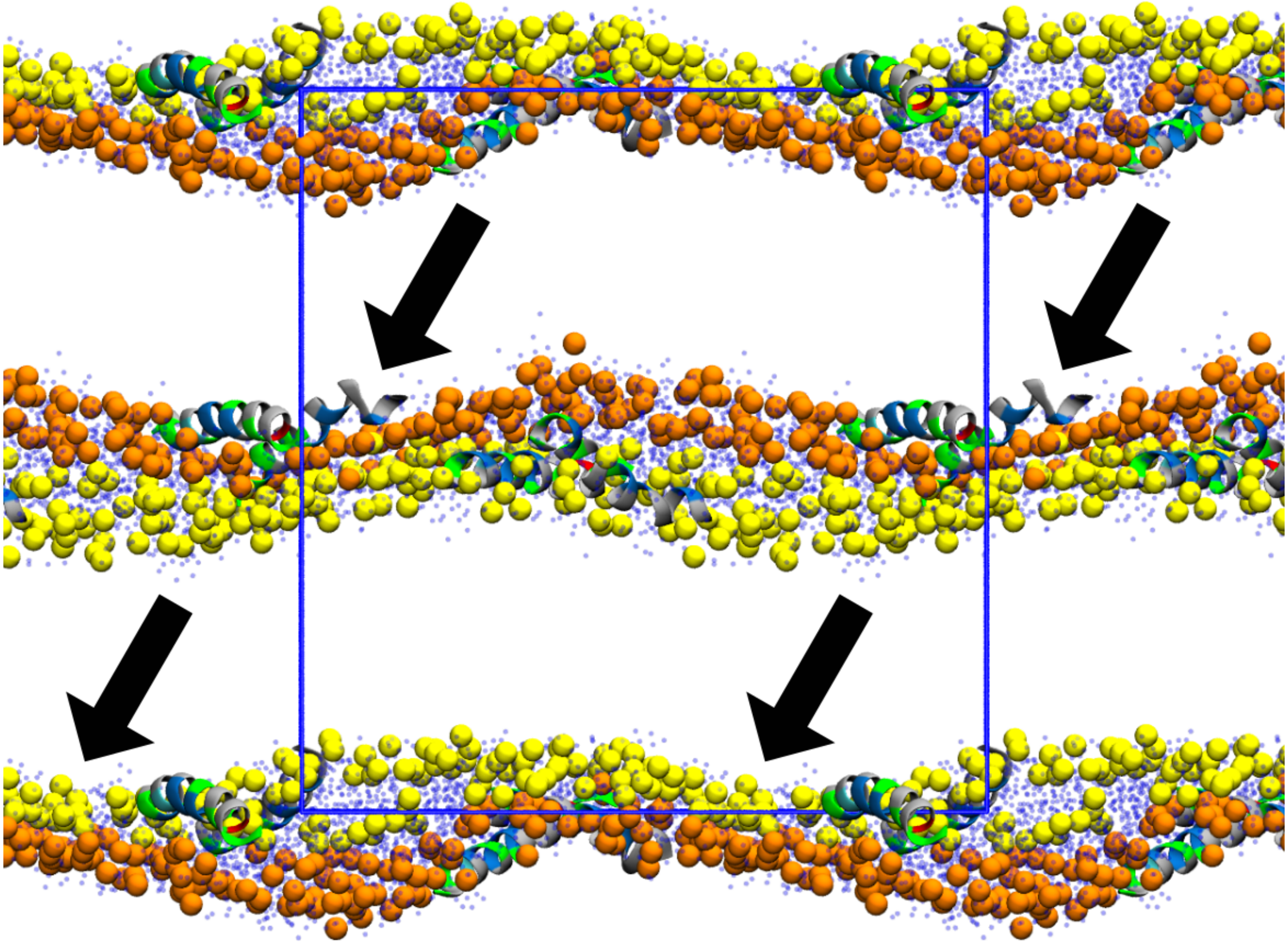

Figure S11: Initial configuration for an all-atom simulation of two lipid bilayers. Blue rectangle marks the simulation cell. Lipid phosphate atoms of first and second bilayers are shown as orange and yellow spheres, respectively. Lipid tails are not shown for clarity. The peptides' secondary structures are displayed in cartoon representation and colored by residue type. Nonpolar: gray, polar: green, acidic: red, and basic: blue. Water oxygens are shown as semi-transparent dark-blue dots.

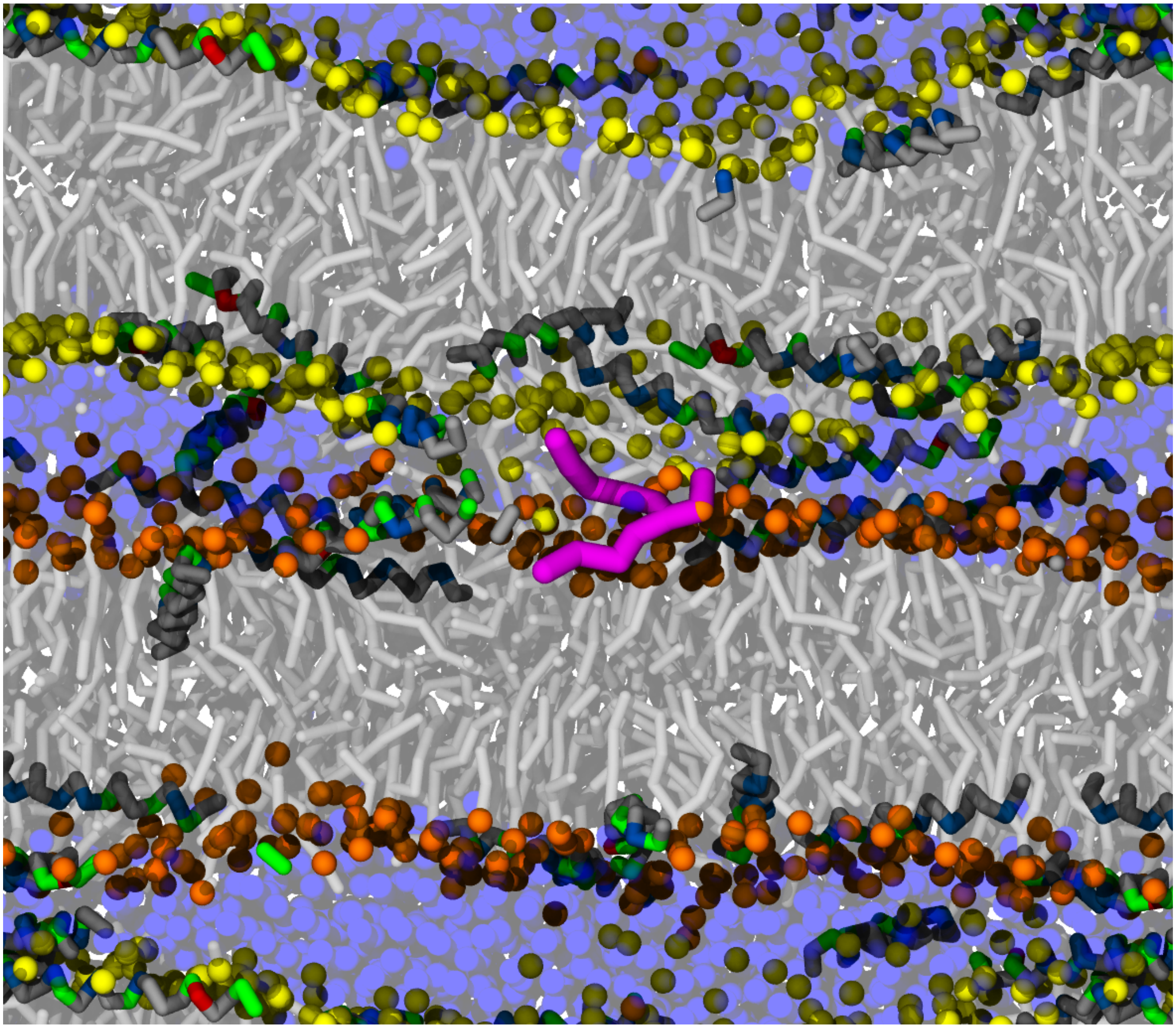

Figure S12: Simulation snapshot of two lipid bilayers connected with a fusion stalk and peptides at P/L = 1:21. Lipid phosphate atoms of first and second bilayers are shown as orange and yellow spheres, respectively. Lipids are shown as light-gray sticks. One lipid connecting both bilayers is highlighted by purple color. Nonpolar: gray, polar: green, acidic: red, and basic: blue. Water beads are shown as semi-transparent dark-blue spheres.

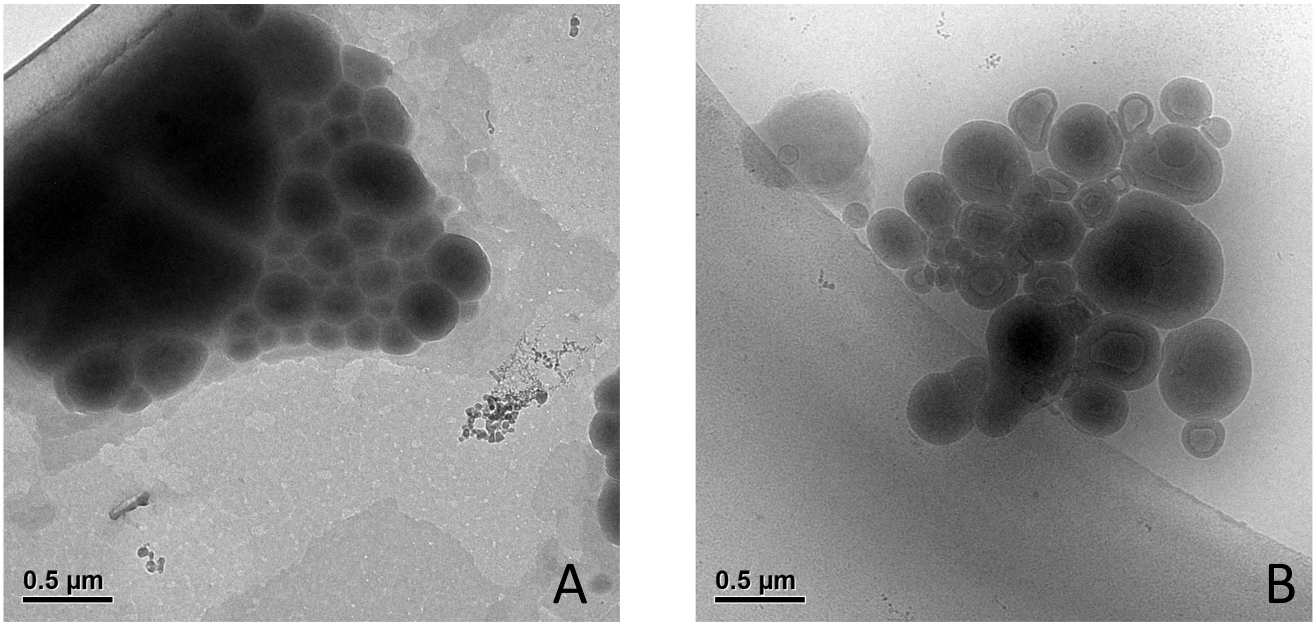


Figure S13: Cryo-TEM images of large unresolved aggregates found for POPE/POPG (3:1) in the presence of the equimolar peptide mixture (panel A) and the hybrid peptide (panel B).
